# Closed-loop feedback control for microfluidic systems through automated capacitive fluid height sensing

**DOI:** 10.1101/221002

**Authors:** L. R. Soenksen, T. Kassis, M. Noh, L.G. Griffith, D.L. Trumper

## Abstract

Precise fluid height sensing in open-channel microfluidics has long been a desirable feature for a wide range of applications. However, performing accurate measurements of the fluid level in small-scale reservoirs (<1mL) has proven to be an elusive goal, especially if direct fluid-sensor contact needs to be avoided. In particular, gravity-driven systems used in several microfluidic applications to establish pressure gradients and impose flow remain open-loop and largely unmonitored due to these sensing limitations. Here we present an optimized self-shielded coplanar capacitive sensor design and automated control system to provide submillimeter fluid-height resolution (~250 μm) and control of small-scale open reservoirs without the need for direct fluid contact. Results from testing and validation of our optimized sensor and system also suggest that accurate fluid height information can be used to robustly characterize, calibrate and dynamically control a range of microfluidic systems with complex pumping mechanisms, even in cell culture conditions. Capacitive sensing technology provides a scalable and cost-effective way to enable continuous monitoring and closed-loop feedback control of fluid volumes in small-scale gravity-dominated wells in a variety of microfluidic applications.

## I. Introduction

Devices for precise fluid handling at the micro-and mesoscale, with characteristic lengths (λ) < 10 cm, are becoming increasingly common across a variety of fields, including biological, pharmaceutical and medical research.^1^ Most microfluidic systems require at least one pump to produce flow, as well as open-or closed-channels or reservoirs to supply, store and collect fluid. Such devices can be categorized in terms of their primary pumping action as mechanical, chemical, surface-, force-field and mixed systems.^1,2,3^ Examples of mechanical pumping at these scales include syringe-pumps, microelectromechanical (MEM) pumps^4^, magnetically actuated pumps^5,6^, vacuum or pressure-driven pumps, peristaltic pumps and centrifugal pumps.^1,7,8^ Chemically-induced flow can be seen in osmotic and effervescence pumps, as well as systems using electrochemical reactions.^1,9,10,11^ Surface-effect pumps include all devices making use of spontaneous capillary forces and substrate wicking to passively induce fluid flow.^12^ Further, force-field pumps are usually defined as non-mechanical including electro-osmotic^3,13^, electrowetting^14^, poroelastic^15^, electrodynamic, thermocapillary and gravity-driven pumps.^1,3,16,17^

Common metrics for performance comparison in microfluidic applications include: operational fluid volumes, flow rates, directionality, persistence time, flow pattern control, fluid capacity, recirculation, optical accessibility, power requirements, heat transfer, and cost of implementation.^1^ Furthermore, criteria such as mechanical, chemical and electrical gentleness during cell or particle handling, as well as material/surface stability against fouling or analyte absorption may also be important to consider.^1^ Understanding the expected performance and inherent limitations associated with each fluid handling and pumping scheme is essential. For example, mechanical pumps connected to open wells or channels do not usually deliver robust steady-state flows for long periods of time as they are bounded by a specific input volume (e.g. syringe pumps), rely on pulsatile flow (e.g. peristaltic, pneumatic and MEM-based pumps)^11,12,18,19,20,21^, are transient in nature (e.g. gravity-driven^22^, capillary-driven^12^ and centrifugal pumps^23^), or are prone to performance variation (e.g. fabrication errors, use-induced stress). Thus, it remains desirable to identify novel approaches to provide robust, scalable and cost-effective monitoring and control of both pumping performance and fluid volume allocation in microfluidic devices.

Among the most attractive targets for improved microfluidic pumping control and volume tracking, we find a variety of open-channel systems such as multi-well cell-culture plates and organ-on-chip platforms^24,25^. In these systems, open wells are usually intended to maintain constant fluid volumes while driven by passive or active recirculation and media oxygenation circuits that extract and replenish fluid from the system.^25^ Variations in input and output pumping performance, as well as unpredictable variations in evaporation rates often lead to significant errors in desired fluid heights over relevant experimental timescales. In the particular case of cell culture applications where high flow-rates are required, fluid height errors in the order of millimeters may render otherwise healthy cultures nonviable after just a few hours of operation.^26^ Moreover, if testing for drug toxicity and other effects related to compound concentration in such experiments, tight fluid volume tracking becomes an absolutely necessity for analysis, which is something largely absent in open-channel fluidic devices used for this purpose.^27^ Indeed, accurate fluid level monitoring capabilities at these scales has been a challenging goal, with current solutions being mostly incompatible with sterile or contact-sensitive applications, or requiring bulky and expensive instrumentation to achieve.^28^ This situation has not only limited the reproducibility of experiments performed in a variety of microfluidic platforms, but has also hindered the use of feedback control to improve the robustness of these systems. Here we present an optimized non-contact capacitive fluid level-sensing technology with submillimeter resolution to provide closed-loop control of a gravity-driven pump for cell culture as a proof-of-concept that demonstrates its use in a range of microfluidic systems.

## II. Materials and Methods

### II.a Feedback-controlled gravity-driven pump setup

A schematic of the assembled testing setup developed to demonstrate the capabilities of our capacitive fluid-level sensor can be seen in Figure 1A. The elements depicted in this system include a capacitive-fluid sensor, a hydrostatic chamber, a microcontroller unit, a bidirectional pump, a microfluidic device and a recirculation container, as well as tubing and connectors.

**Figure 1.**
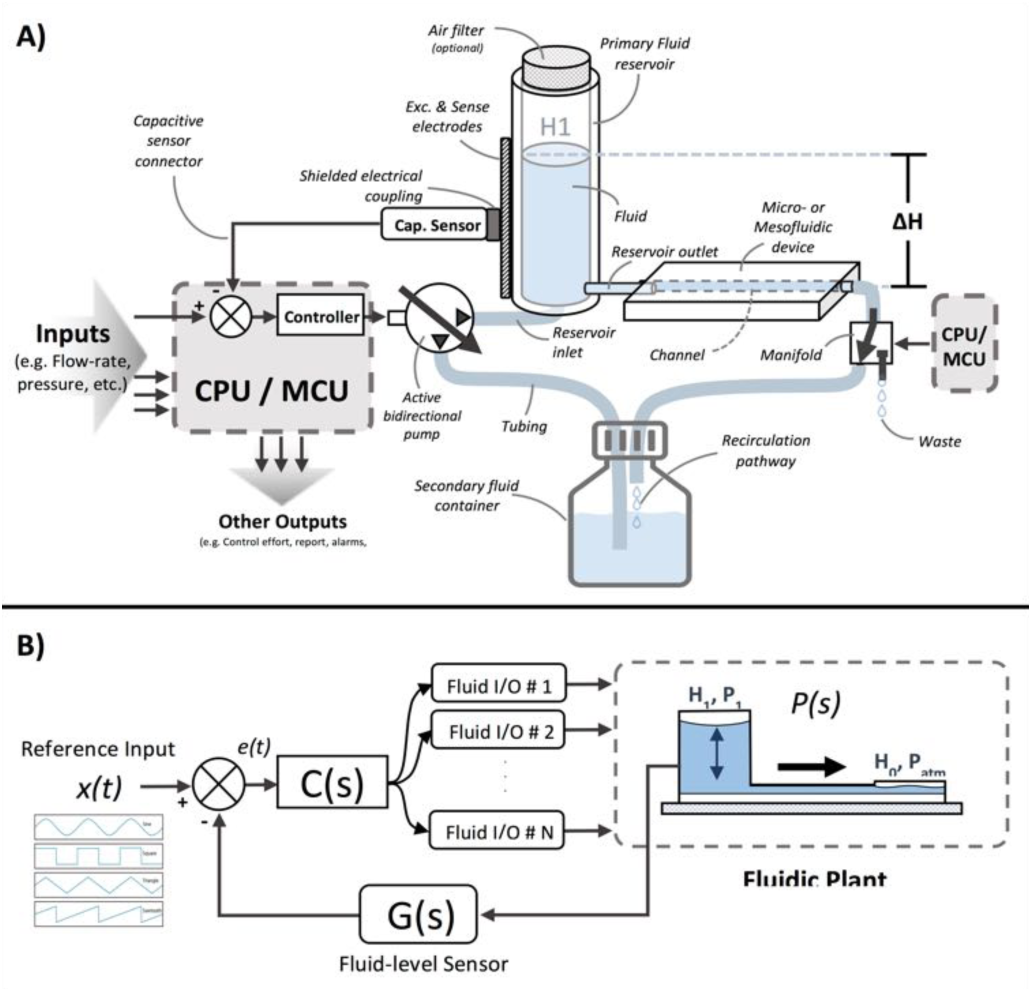
A) Schematic of the proposed closed-loop controlled gravity-driven microfluidic setup with capacitive fluid-level sensing. In this system, the target microfluidic chip is driven by only one hydrostatic pressure chamber, although multiple chambers could be used. B) Simplified closed-loop feedback-control block diagram of the gravity-driven setup. Reference input x(t) is user-defined and compared to the measured state (i.e. fluid height, pressure) to calculate the error e(t). Fluid I/O’s refer to the active bidirectional supply pumps that control the height in the primary chamber. C(s), P(s) and G(s) are the controller, fluidic plant and sensor transfer functions respectively.

In this testing setup, the fluid level in the primary fluid reservoir is set to the desired position (H1) to act as a hydrostatic chamber, which drives gravity-driven flow through a meso-and/or microfluidic chip. This chip is located inside a device holder to be connected to the fluidic circuit via standard tubing and microfluidic connectors. The height difference (ΔH) with respect to the chamber outlet produces a hydrostatic pressure (*P*_*in*_) at the inlet of the connected microfluidic channel such that *P*_*in*_=ρgΔH; where ρ≈1 g/cm^3^ (fluid density for water) and g≈9.81 m/s^2^. The flow rate imposed through the microfluidic device (Q) is then related to the pressure gradient between the inlet and outlet of the microfluidic channel (Δ*P*) as well as its fluidic resistance (R) such that Q= Δ*P/R*. During operation, the fluid level in this primary reservoir is continuously monitored using the non-contact fluid-level sensor. The recorded fluid level is then fed to a microcontroller unit (MCU) implementing a closed-loop algorithm (Figure 1B). This system takes a user-defined reference input signal *x(t)* (i.e. height, pressure, flow rate) and compares it to the measured state to produce an error signal *e(t)*. The generated error feeds a control law that actuates a bidirectional pump capable of actively increasing or decreasing the fluid height in the primary fluid reservoir to track the reference input *x(t)*. Standard tubing, connectors and a secondary container complete the fluidic circuit, all of which allow for stand-alone operation. A prototype of this system, including required electronic components and other structural elements (e.g. controller box, sensor/chamber holder, microfluidic device holder) can be seen in Figure 2A. More detail on the design and fabrication of each of the components and modules is presented in further sections.

**Figure 2.**
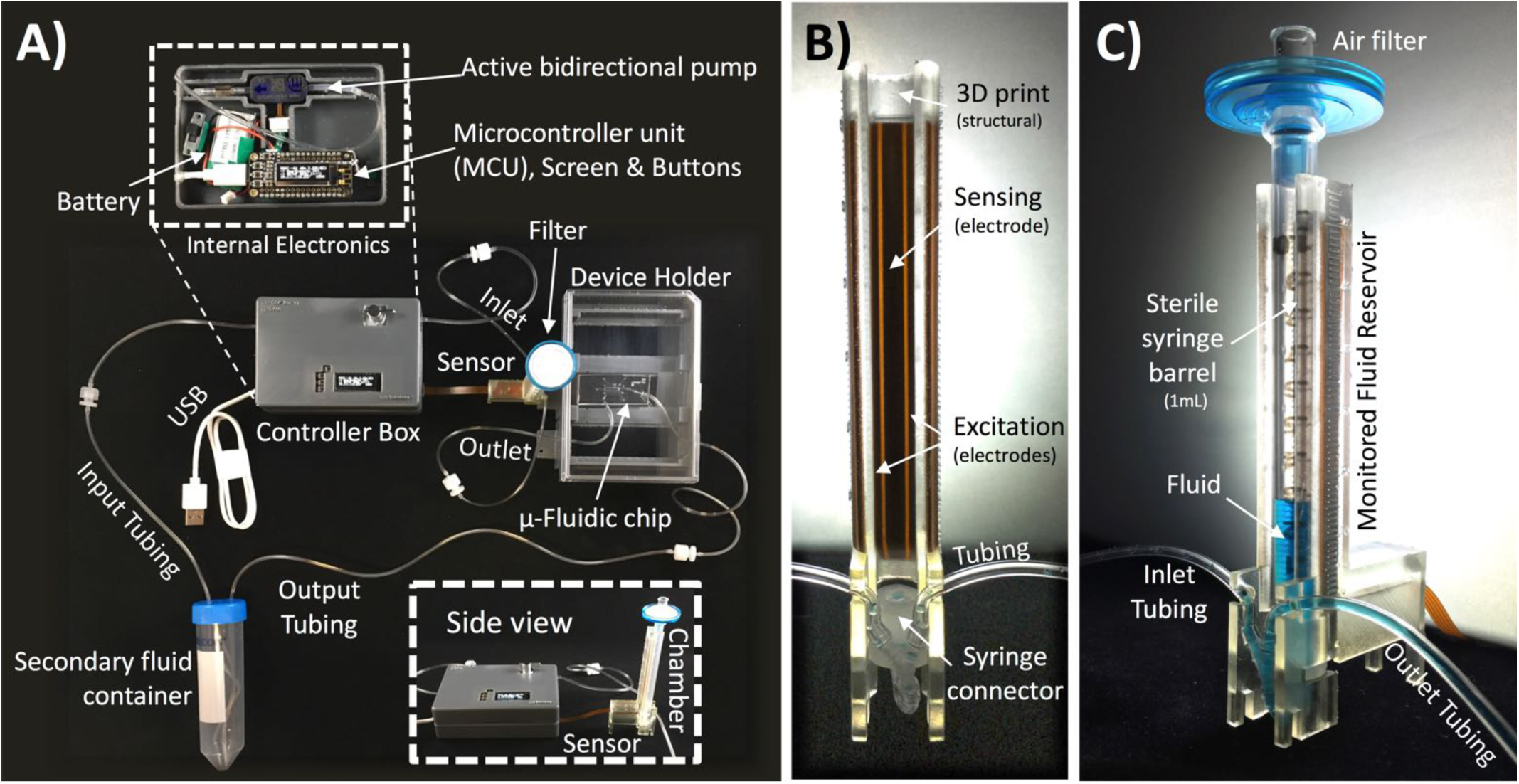
A) Fabricated testing setup used to perform the validation experiments of height-based closed-loop control of open-well microfluidic systems. B) Detail of embedded non-contact capacitive fluid-level sensor with custom-designed electrodes. C) Assembled gravity driven reservoir with embedded capacitive sensing and a single upper air filter (0.22 μm pore size) for sterile use in cell culture applications.

#### II.a.1 Hydrostatic fluid chamber

Figure 2C shows the assembled hydrostatic chamber with capacitive sensing used in the feedback-controlled gravity-driven setup. In this assembly, a sterile 1mL Plastipak™ graduated syringe barrel (Becton Dickinson, Rutherford, USA) is oriented vertically and connected to a 1/16” polypropylene barbed quick-turn coupling socket (McMaster Carr, Robbinsville, USA) within a 3D printed structure to form a biocompatible and sterile reservoir with dimensions (H_total_=60 mm, ID=4.78 mm). A 0.22 μm pore Fisherbrand™ filter (Thermo Fisher Scientific, Waltham, USA) is placed on top of the assembly to allow for air flow while maintaining sterility. This hydrostatic chamber is connected to the pumping and recirculation circuits through 1/16” ID polypropylene tubing and a compatible nylon tube-to-tube wye connector (McMaster Carr, Robbinsville, USA). Gravity-driven flow is determined by the height of the fluid column and the downstream resistance of the system. The microfluidic device is connected using standard tubing and located within the device holder (Figure 2A) to reliably control its vertical position with respect to the bottom of the fluid column. The non-contact capacitive fluid-level sensor is in close proximity to the fluid (^~^2mm) in the 3D printed structure (Figure 2B) so as to monitor fluid height. Additional details on the components used for the assembly of this monitored hydrostatic chamber can be found in Figure S1 of the supplemental material.

#### II.a.2 Capacitive fluid-level sensor for microfluidics

Our optimized capacitive fluid-level sensor for microfluidic applications consists of self-shielded coplanar electrodes connected to an AD7746 24-Bit Σ-Δ capacitance-to-digital converter in differential mode (Analog Devices, Norwood, USA). Figure 3A illustrates the layout of this sensor in conjunction with the monitored fluid reservoir. In our design, two pairs of excitation electrodes (diagonal pattern in Figures 3A and 3B) are positioned around two sensing electrodes (dotted pattern) to measure fringing capacitance in the direction of the fluid as shown in Figure 3C. We refer to this electrode design as excitation-sensing-excitation/inter-digitated arrangement (ESE-ID). Two symmetrical gaps separate the excitation electrodes from the central sensing electrodes. The gap length is a parameter that is affects the penetration depth of the most sensitive fringing pathways in other types of capacitive sensors using coplanar electrode arrangements.^29^ Thus, L_gap_ can be iteratively adjusted to achieve optimal sensing in other applications with different fluid-sensor wall thicknesses. For our hydrostatic chamber prototype in Figure 2C, L_gap_=0.75mm was heuristically determined (from several design iterations) as sufficiently small to allow for capacitive sensing using our electrode geometry.

**Figure 3.**
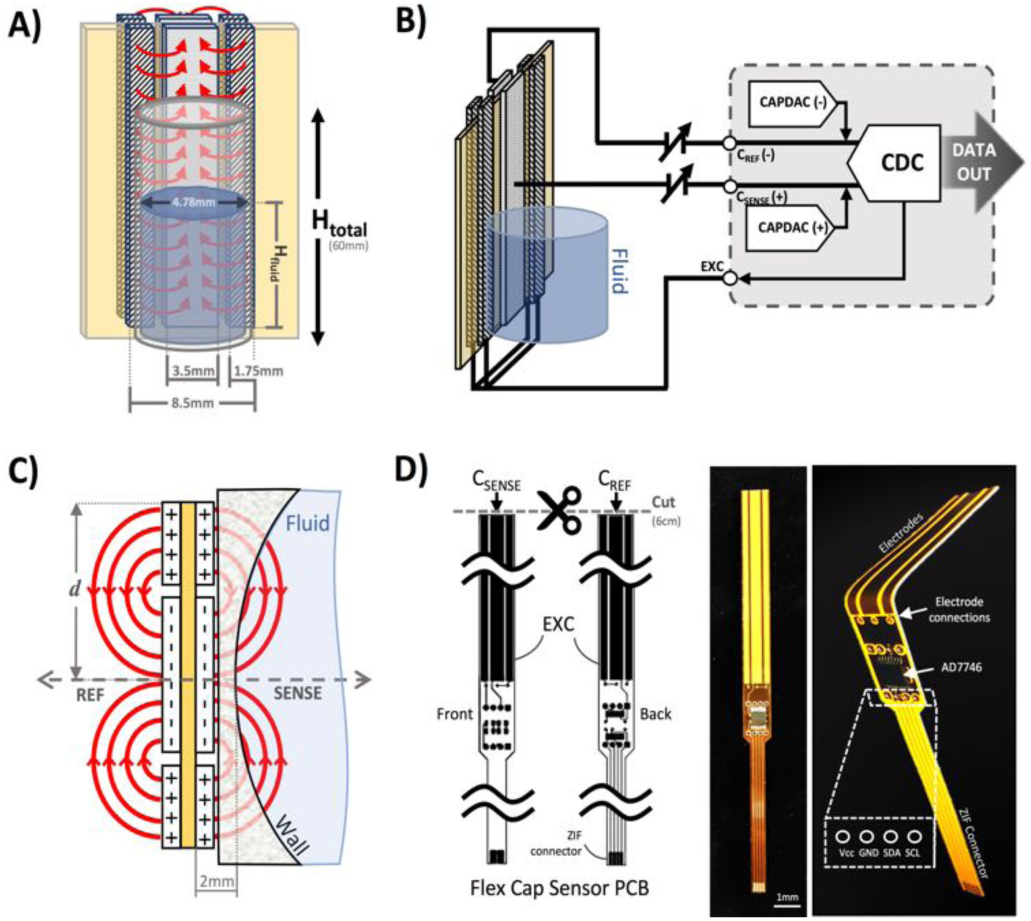
Design of the capacitive fluid-level sensor for microfluidic applications. A) Planes filled with diagonal patterns are excitation electrodes, while planes with dotted patterns are sensing and reference electrodes. The yellow plane is a middle polyimide dielectric layer, while the blue cylinder represents the monitored fluid reservoir in experimental setup. B) Shows the sensing, reference and excitation electrodes connected to their respective terminals on the CDC chip. C) Cross-sectional detail of charge distribution and fringing capacitance fields (red arrows) among sensing, reference and excitation electrodes. The relative position of the fluid reservoir wall and the sensing electrode is also shown. D) Flex-PCB layout and images of fabricated sensor. **CDC**=Capacitance-to-digital converter; **CAPDAC**=Programmable on-chip digital-to-capacitance converter; **GND** = Ground supply; **PCB**= Printed circuit board; **SDA/SCL**= I^2^C communication pins; **Vcc**=Power supply.

The AD7746 chip is connected to the front sensing electrode via the C_sense_(+) terminal, while the back sensing electrode is connected to the reference terminal C_ref_(-) to allow for differential measurements. The excitation electrodes of both sensing and reference planes are connected to the same excitation (EXC) terminal. The differential mode of the AD7746 chip was selected to maximize the robustness of capacitance measurements, while the addition of a symmetrical reference arrangement at the back of the sensing layout was designed to maximize signal-to-noise ratio. This effect can be explained by referring to the expected fringing capacitance pathways on the presented mirrored electrode design (Figure 3A), which shows a self-shielding effect within the sensor.

In commonly used capacitive sensing instruments, the region separating the electrodes from a target fluid is usually made of materials with low electrical permittivity (ε) such as plastic, glass or air.^30^ This renders the capacitance due to the plastic and air small as along as ε_Air <_ ε_Plastic ≪_ ε_Fluid_ is maintained. Thus, the capacitance attributable to the region next to the fluid can be approximated to the total capacitance detected at the C_sense_(+) terminal. Furthermore, in previously described coplanar sensor designs^30^, the reference electrodes are usually situated in the same plane as the main sensing electrode (next to a region constantly filled with fluid).

This traditional configuration simplifies the sensor compensation for different kinds of fluids and temperature changes, but it also brings several limitations in terms of footprint, minimum detectable fluid volume and achievable signal-to-noise ratio. Our sensor prototype is implemented using a 3-layer flexible printed circuit board (Flex-PCB) with total thickness of 0.2mm. Electrodes are defined as 0.5 oz copper layers with 17.5μm thickness, while a 55 μm polyimide film was used as dielectric. A layer of dielectric film lies between the mirrored electrodes, as well as at the top and bottom of the sensor to protect the conductive material from corrosion (Figure 3D). Connection traces between sensing circuit and electrodes use 0.127 mm traces. The electrodes are made with an original length L_total_ = 20 cm, and then are cut at the top end to fit the 6 cm fluid reservoir. This design allows for use of the sensor in longer hydrostatic chambers with minimum modification of the sensor design. The separation of the mirrored reference electrode from the main sensing plane was achieved using a 75 μm adhesive-polyimide-adhesive dielectric layer. The charge distribution imposed by this design directs the fringing fields of the sensing electrode plane preferably towards the fluid, while directing the fields of the reference electrode plane towards the back of the sensor. This configuration is more compact than traditional coplanar capacitive sensors, and enables the detection of smaller increases in fluid height and more effective compensation of external parasitic capacitances (e.g. user’s movement). For this configuration, the capacitance associated with the sensing and reference coplanar electrodes can be approximated as^31^:

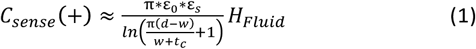

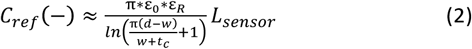

where C_*sense*_(+) and C_*ref*_(-) are the capacitances of the sensing and reference electrodes respectively, *d* is the average diameter of the fringing arcs between the sensing and the excitation electrodes. *L*_*sensor*_ is the total length of the sensor, *H*_*Fluid*_ is the fluid height, *t*_*c*_ is the thickness of the electrode conductor (assumed to be constant across all electrodes), and ε_O_ is the permittivity of free space (ε_O_ ≈ 8.9 × 10^−12^ F/m). Due to symmetry, equations (1) and (2) are only valid for the case in which the width of the sensing electrode (*w*) is approximately equal to the added width of both excitation electrodes. The relative permittivity associated with the sensing region (ε_*S*_) and the reference region (ε_*R*_) in equations (1) and (2) can be determined by examining the ratio between the average diameter of fringing arcs (*d*) and the thickness of the material separating the fluid from the conductor (*t*_*w*_).^32^ The thickness *t*_*w*_ considers the flex-PCB dielectric and plastic wall and is only used to determine ε_*s*_ and ε_*R*_ according to the following rules: For *d/t*_*w*_≫1, ε_*s*_ = ε_*R*_ ≈ 1; whereas for *d/t*_*w*_ ≈ 1, ε_*s*_ = (1+ ε_*fluid*_)/2 ≈ 40 and ε_*R*_= (1+ ε_*plastic*_)/2 ≈ 1. All previous approximations assume a relative permittivity for the fluid (ε_*fluid*_) around 80 at 20 °C under a 1 kHz excitation, while ε_*plastic*_≈2 for a bulk plastic dielectric material. Since our testing setup has *d*=2.5 mm and *t*_*w*_=2 mm, it follows that *d/t*_*w*_=1.25 ≈ 1 confirming that the second ratio condition applies. Dividing C_*ref*_(-) from C_*sense*_(+) and reordering terms we can approximate the fluid level in the reservoir as follows^32^:

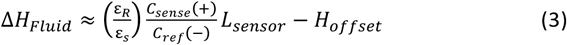

where ε_*R*_/ε_*S*_ = (1+ ε_*plastic*_)/(1+ ε_*fluid*_) is just the proportionality constant in equation (3), which depends on the target fluid and the bulk dielectric material. This constant can be calculated for compensation purposes by performing a single capacitance measurement at a known fluid height. Finally, an additional compensation offset (*H*_*offset*_) may be used to correct for mechanical inaccuracy due to sensor placement and to adjust for absolute height according to a common height reference as seen in equation (3). The design and fabrication files for the implemented sensor are available for download in the supplemental material.

#### II.a.3 Electronic feedback-control hardware

After acquisition of capacitance readings by the AD7746 chip, the digitized measurements are transmitted to a microcontroller board HUZZAH^®^ ESP8266 Feather (Adafruit Industries, New York, USA) via a ZIF connector and I^2^C serial communication terminals to perform feedback control computations. This board was selected as our control hardware due to its low-cost (<$20 USD), ease of programming (using Arduino^®^ syntax), availability of open-source design files, power efficiency and integrated wireless capabilities. This microcontroller board was then connected to a stackable custom-made active pumping board, and a FeatherWing OLED I/O interface board (Adafruit Industries, New York, USA) with assembled push buttons and a screen for offline parameter visualization. The detail in Figure 1A shows the assembled control hardware, while additional images of the individual components of the controller can be found in Figure S2 from the supplemental material.

#### II.a.4 Bidirectional supply pumps

In order to actively control fluid height within the hydrostatic fluid chamber, two Bartels Microtechnik mp6 piezoelectric pumps (Servoflo, Lexington, USA) are driven in a bidirectional configuration by the pumping board using two commercially available mp6-OEM driver circuits (Servoflo, Lexington, USA).^33^ The flow-rate of these piezoelectric pumps is controlled using a pulse-width modulated (PWM) signal generated by two independent output channels from the HUZZAH^®^ ESP8266 Feather. This type of pump was selected due to its small size, fast response, high dynamic range, and chemical inertness. Other active bidirectional pumps may be used as long as their response time is faster than the characteristic time constant of the fluidic plant to be controlled. All PCB fabrication files to reproduce the pump board are included in the supplemental material.

#### II.a.5 Control and user-interface software

The AD7746 acquisition routine and a proportional-integral (PI) control law were implemented using the Arduino^®^ in-system programmer (ISP) on the ESP8266 microcontroller unit. We also used the Arduino and Adafruit ESP8266 libraries to facilitate integration of this platform. Capacitance measurements were obtained every 10 milliseconds using a timer-driven interrupt. A sensor calibration routine was also implemented, so as to allow the user to record a base capacitance offset as well as the calculation of the fluid-dependent proportionality constant (ε_*R*_/ε_*S*_). Routines to follow constant and pre-programmed dynamic fluid height profiles were also implemented in non-volatile memory of the ESP8266 (EEPROM). Acquired data and system control parameters were transmitted via USB and wirelessly via Wi-Fi to a laptop and then converted to CSV format for analysis. The code used for the conducted experiments is included in the supplemental material.

#### II.a.6 Microfluidic devices and fluidic circuit

Two simple microfluidic devices were used to test usability and validate functionality of our feedback-controlled gravity-driven setup. These microfluidic devices are: 1) A custom fabricated single channel polydimethylsiloxane (PDMS) chip, and 2) A commercially available chip for cell culture as described later. For most of our tests the output channel of the driven microfluidic device was connected to a dripping recirculation pathway that maintained the outlet of the microfluidic channel at atmospheric pressure as depicted in Figure 3A. It is also possible to use two of these hydrostatic pressure chambers at both ends of the microfluidic chips to allow for bidirectional gravity-driven flow, as well as regulating the device average pressure.

#### II.a.7 Structural and other components

Structural elements such as the sensor/chamber holder, controller box and microfluidic device holder were fabricated from stereolithographic (SLA) resin using a Form 2 3D printer (Formlabs, Somerville, USA). The hydrostatic chamber and sensor holder are made from clear resin to allow for direct optical visualization of the monitored fluid front for validation purposes. The design and fabrication files of these structural components are also included in the supplemental materials.

### II.b System testing and validation

#### II.b.1 Validation of capacitance measurements

In order to characterize the accuracy of the capacitance measurements obtained by our sensor, we first isolated the AD7746 circuit of the Flex-PCB. We then placed capacitors of known values between the sensor’s C_sense_(+) and EXC terminals to record their values. The tested capacitances were verified using a calibrated E4981A capacitance Meter (Keysight Technologies, Santa Rosa, USA) for comparison purposes. The results of this test are shown in Figure 4A.

**Figure 4.**
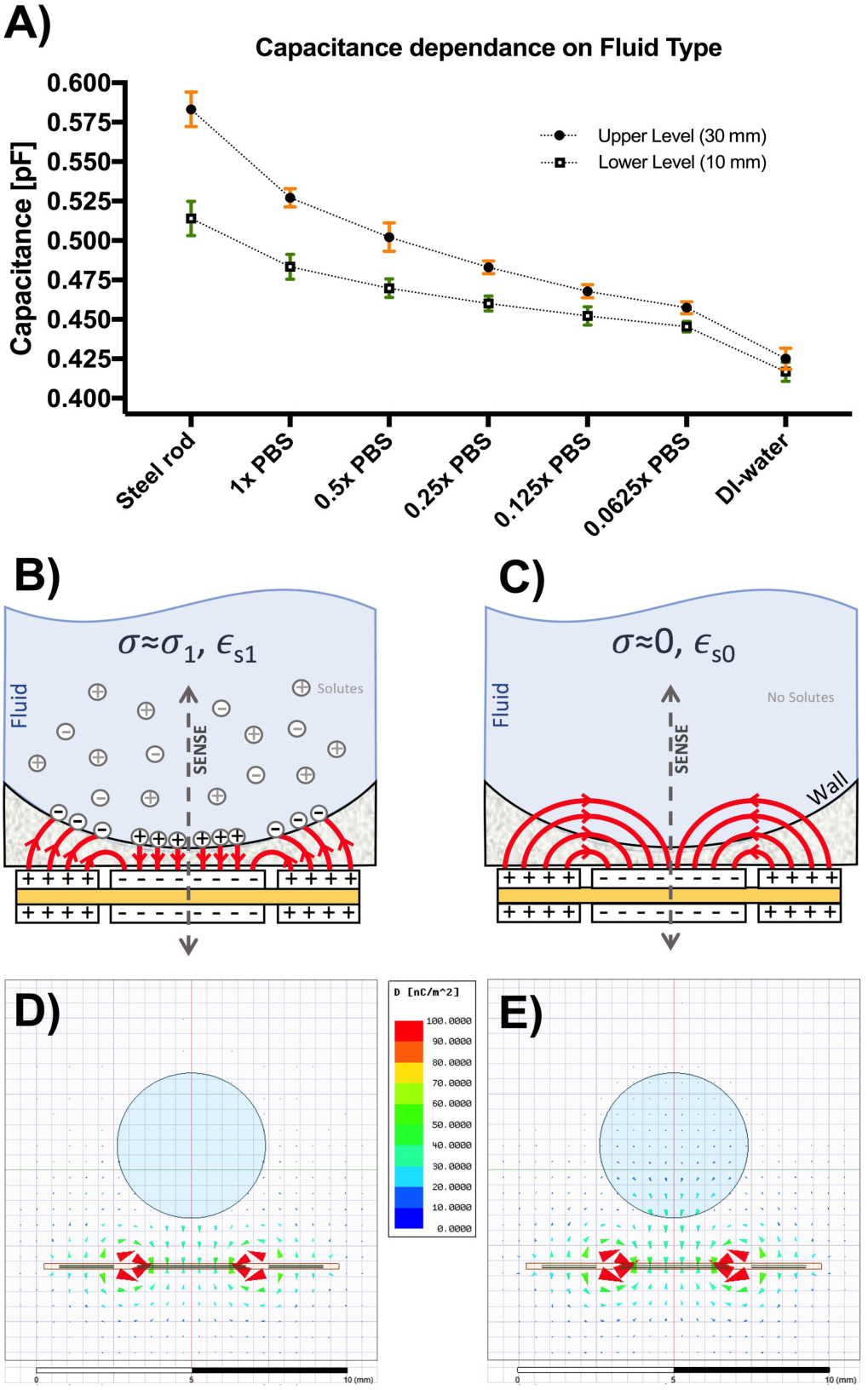
A) Results on dependence of capacitive sensing on fluid type. Use of fluids with high concentration of solutes forming free ions lead to higher capacitance measurements and ranges. Error bars representing standard deviation of upper measurements (h=30mm) are shown in orange, while error bars of the lower measurements (h=10mm) are shown in green. B) Capacitive fringing model for electrolytic fluids with free ions. C) Capacitive fringing model for non-conductive fluids with high dielectric constant. Fringes at the fluid-wall interface exhibit a change in angle towards the symmetry axis (not shown) depending on properties of wall material and fluid. D) Finite element analysis (FEA) of the electric displacement flux density for 1xPBS target fluid. E) FEA simulation of electric flux density for DI-water as target. In both D) and C) the blue circle is the target liquid, and the outer box is plastic material (ε_r_ = 2.7, σ = 0 S/m) following the same dimensions as the physical prototype chamber. Outside the plastic box region there is air. The thin orange contour surrounding the sensor electrodes is a polyamide film (ε_r_ = 4.3, σ = 0 S/m). The color of the vectors represents the magnitude of the flux density *D* (nC/m^2^) shown in the middle color bar. The size of the arrows is proportional to the magnitude of the flux density to facilitate interpretation.

#### II.b.2 Basic fluid-height tracking

After characterization of the sensing circuit, a basic fluid height “challenge” within an isolated hydrostatic chamber with an embedded capacitive sensor was performed in triplicate using a 1x phosphate buffered saline (PBS) solution. The fluid level in this reservoir was controlled using a calibrated syringe pump 11-Pico PLUS Elite (Harvard Apparatus, Holliston, USA). This pump was programmed to produce a dynamic oscillating change in fluid volume over the reservoir’s entire height range (6 cm) during a 20 s period. Readings from our capacitive sensor were taken every 10 ms (without averaging) and compared to the fluid volume supplied by the syringe pump. Fluid height in the chamber was also verified using video recordings and image processing to track the fluid-air interface visualized through the translucent regions of the fluid reservoir. The results are shown in Figure 4B.

#### II.b.3 Gravity-Driven Flow Rate Validation

To validate flow rates established using a given hydrostatic pressure (i.e. fluid height), we connected our pump to a 50 cm long silicone tube with inner diameter of 1/32” and measured the mass of 1x PBS displaced over a one minute duration (n=3) using a laboratory grade scale. The Poiseuille equation was used to calculate the expected flow rate given a hydrostatic pressure difference ΔP=8μLQ/(πR^4^), where ΔP is the hydrostatic pressure difference, L is the length of the tubing, μ is the dynamic viscosity of PBS which is assumed to be close to that of water (8.9x10^−4^ Pa.s at 25 °C), Q is the volumetric flow rate and R is the inner radius of the tubing. Additionally, the hydrostatic pressure was calculated as ΔP= ρgh, where ρ is the density of PBS which is approximately that of water (1 kg/m^3^), g is gravity and h is the total fluid height above the outlet feeding to the collection tubes used for weight measurement. In order to achieve high flow rates, we added an additional 53 mm offset to the fluid column by placing the collection tubes below the gravity driven pump outlet. The obtained flow-rate measurements (in triplicate) were compared to expected values using Poiseuille’s equation as shown in Figure 4C.

#### II.b.4 Characterization of fluid-type dependency

As described earlier, the gain of our capacitive sensor design is expected to change depending on the charge distribution resulting from the conductivity and the electrical permittivity of the target fluid. Therefore, changes in charged solute concentration can affect these readings significantly. To characterize such effect, we conducted a titration experiment in triplicates using 1x, 0.5x, 0.25x, 0.125x and 0.0625x PBS, with deionized (DI) water and a steel inserts of known lengths as controls. Two inserts were made by cutting 10-and 30-mm sections from a tight-tolerance multipurpose O1 tool steel rod with 0.1750” diameter (McMaster Carr, Robbinsville, USA). The associated capacitance value from all fluid conditions and materials was recorded at two fixed fluid heights: lower Bound (H_0_=10 mm) and upper bound (H_1_=30 mm) as shown in Figure 5A. To confirm experimental results, we also computed the electric displacement flux density (*D*) around relevant target fluids (i.e. 1xPBS solution, DI-water), via finite element analysis (FEA) using the AC conduction field solver of ANSYS Maxwell software (ANSYS, Inc., Canonsburg, USA) as shown in Figures 5D and 5E. In these simulation, the two middle electrodes were fixed to zero voltage, and the four electrodes on the side were excited with a sinusoidal voltage of 5V amplitude at an excitation frequency f_EXC_ = 32 kHz. These conditions are analogous to the settings used by the AD7746 24-Bit Σ-Δ capacitance-to-digital converter in the developed sensor.

**Figure 5.**
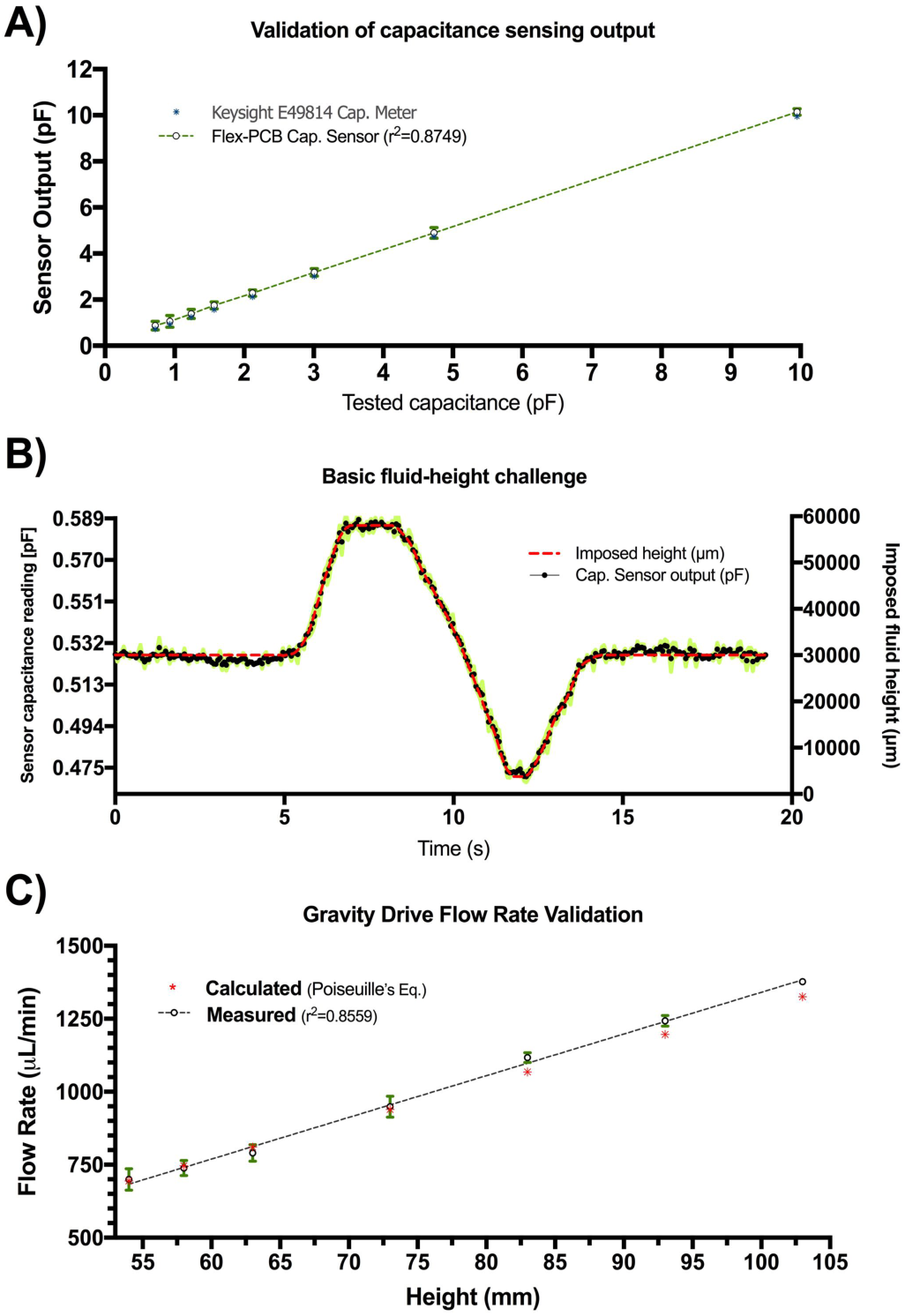
A) Results of capacitive sensing circuit validation experiments. Capacitances measurements with the developed sensing circuit matched real values within <0.1pF error. Circles denote average of experiment conducted in triplicate at a single capacitance value with the error bars shown in green. A linear fit of the measured values is also shown. B) Results of basic fluid-level tracking challenge using capacitive sensing. The dashed profile (red) is the expected fluid-height as imposed by the calibrated syringe pump. Black points refer to the averaged capacitive sensor output for the three replicates at each time point. C) Measured and calculated flow rates as a function of total fluid height in gravity-driven pump. The measured values (circles) with green error bars, closely approximate the theoretical values calculated using Poiseuille’s equation (asterisks). A linear fit of the measured values is also shown.

#### II.b.5 Flow-rate tracking and calibration of external pneumatic micropumps

Since accurate contactless assessment of flow in open-channel microfluidics is a valuable potential application for the present technology, we conducted a series of proof-of-concept experiments to demonstrate flow-rate extraction and open-loop pumping calibration based on continuous fluid-level monitoring. The same previously described fluid-height tracking methodology was used, except that in these experiments, the calibrated syringe pump was programmed to impose specific flow-rate profiles (instead of volumetric changes) ranging from 100 nL/min to 1 mL/min These conditions included constant, ramp and intermittent flow-rates in input and output mode. Fluid-height changes over time were measured in triplicates and compared to theoretical flow estimates using the pre-programmed flow-rate parameters imposed by the calibrated syringe pump. Extraction of flow-rate was approximated by generating a linear fit of fluid volume change over time every 1000 samples (t=10 s). Experiments were limited to a maximum input/output volume (V_max_=0.5 mL) and a maximum experimental period (T < 400s). The results are shown in Figure S3 of the supplemental material.

In a subsequent experiment, this height-based flow-rate tracking methodology was used to characterize and calibrate two presumably identical open-loop pneumatic diaphragm micropumps. These micropumps were set to supply and extract fluid from the same monitored reservoir at 1 μL/s. These pumps were fabricated in acrylic using a CNC mill and a thin polyurethane membrane according to a previously reported protocol.^34^ Both pumps were actuated for 40 min at 1 Hz (stroke volume=1 μL) under 37 °C and 95% humidity to observe fluid-height drift (in triplicate). After confirming adequate operation of both pumps, any observed volume drift in the monitored reservoir is assumed to be caused by small differences in input/output pumping performance attributable to fabrication variations or head pressure effects. After combined drift characterization, flow-rates were independently tracked in triplicate for each pump to recalibrate their actuation frequency and adjust for errors. After calibration, fluid-height drift was again characterized for 40 min and compared against uncalibrated behavior (n=3) as shown in Figure 6.

**Figure 6.**
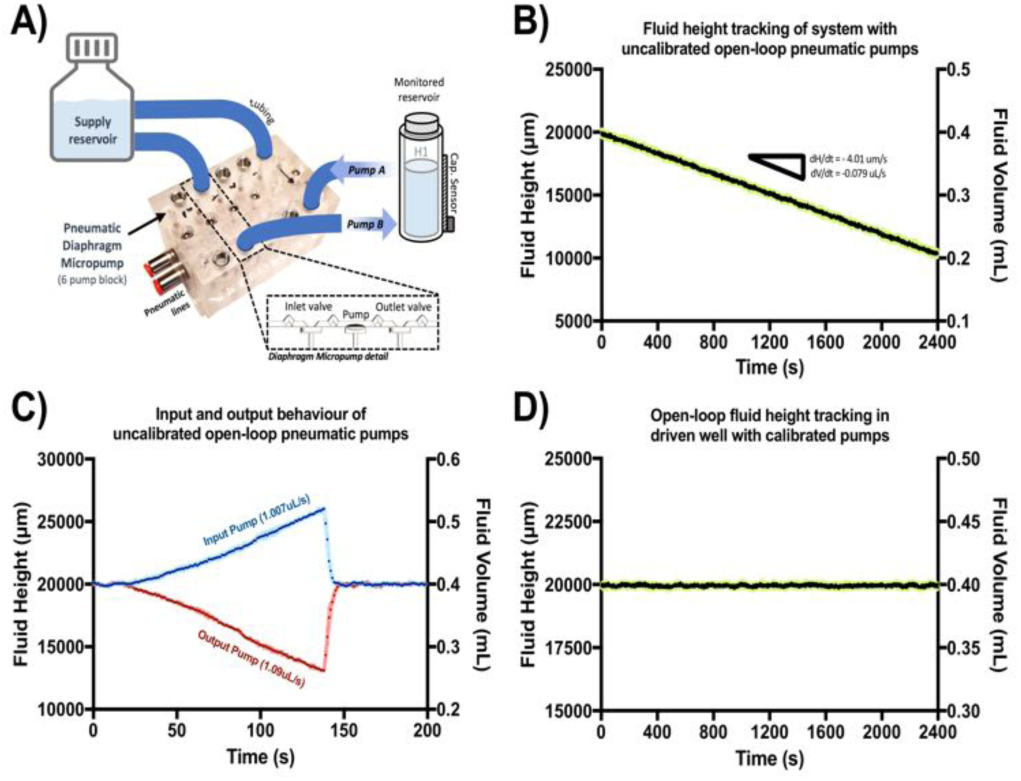
A) Testing block with six parallel pneumatic diaphragm micropumps. Two pumps were used to drive fluid in and out the monitored fluid reservoir. B) Fluid-height change over time (40 min) as measured by the capacitive sensor in the system driven by unbalanced input/output micropumps. C) Overlay of isolated experimental input and output behavior of each pneumatic diaphragm micropump. Blue profile denotes fluid-height over time produced by pump A (input) alone, while red profile shows the same for pump B (output). D) Fluid-height change over time (40 min) in the same system after micropump calibration using extracted flow-rate from capacitive sensing readout. Black points denote average for the three replicates at each time point. Standard deviation is shown in light green.

#### II.b.6 Validation of closed-loop control of gravity driven pump

The performance, robustness and dynamic range of the proposed closed-loop feedback control system was characterized using the entire monitored setup as seen schematically in Figure 1A. A single-channel microfluidic device made in a PDMS (length=50 mm, width=1 mm and height=0.2mm) was connected to the outlet of the hydrostatic chamber with capacitive sensing. After the chip was connected to the setup, sensor calibration and flow testing was performed to assess the emptying time constant of the hydrostatic chamber given the fluidic resistance from the connected microfluidic chip. After this, a closed-loop feedback control mode was activated to automatically control fluid input/output from the secondary piezoelectric pumps to maintain a target height (ΔH=30 mm). For this experiment, the hydrostatic chamber was monitored over a period of 48 hours inside an incubator (at 37 °C, 5% CO_2_ and 95% humidity). Presence of flow through the microfluidic device was confirmed by observing dripping fluid within the secondary recirculation container. This experiment was conducted in triplicate and the fluid height was also verified using video recordings of the fluid front to assess drift as shown in Figure 7A

**Figure 7.**
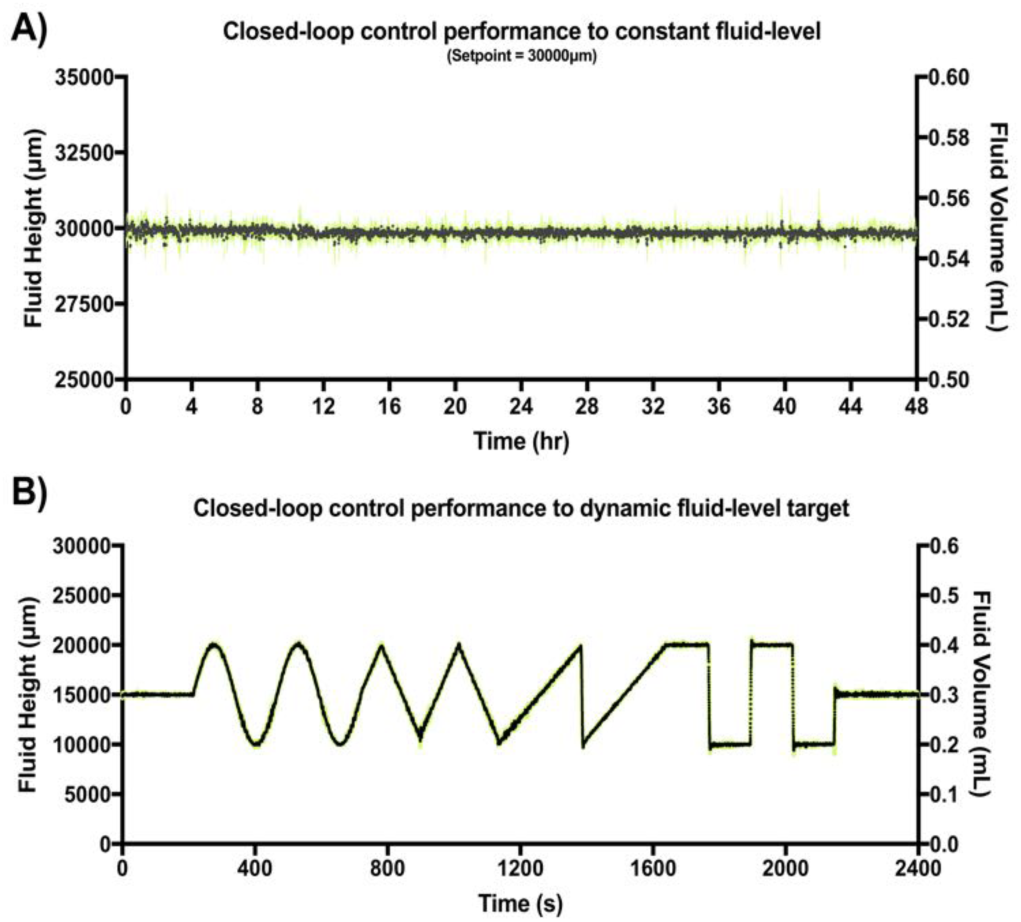
A) Results of closed-loop feedback control for a constant set point (ΔH=30mm). Black points refer to the averaged capacitive sensor output for the three replicates at each time point. Error bars are shown in green. B) Results of closed-loop feedback control for a dynamic set point using constant, sine, triangular, saw tooth and step waveforms. Black points refer to the averaged capacitive sensor output for the three replicates at each time point. Error bars representing standard deviation are shown in green.

After verifying adequate closed-loop performance for a constant fluid-level set-point, a dynamic fluid-level target experiment was conducted under similar experimental conditions. However, in this case the target was pre-programmed to be a dynamic fluid-level profile stored in non-volatile memory of the MCU. This profile was a 40-min sequence including constant, sine, triangular, saw tooth and step waveforms (two periods each). This experiment was also conducted in triplicate within an incubator (37 °C at 95% humidity) using video recordings to verify location of the fluid front. Results are shown in Figure 7A.

#### II.b.7 Cell culture experiments

In order to show biocompatibility of the setup, a 24-hour cell culture experiment was performed using iPSC-derived vascular endothelial cells as an example cell type. In this test, an IBIDI μ-Slide VI 0.4 channel slide (IBIDI, Martinsried, Germany) was coated with human fibronectin (Life Technologies, Woburn, USA) at a concentration of 30 μg/mL for 1 hour at room temperature. Induced Pluripotent Stem Cell (iPSC) derived endothelial cells (CDI, Madison, WI) were seeded in all the channels, allowed to adhere for 3 hours then excess cells were washed away using two successive media changes. Results are shown in Figure 8. The device was cultured for 24 hours in the supplier-recommended media under standard incubator conditions. After assembly, the gravity-driven setup was sterilized by circulating 70% ethanol through the entire fluidic circuit for 5 min. Ethanol was removed by air drying within a sterile hood and then flushing with two cycles of sterile DI water. After sterilization, the seeded IBIDI μ-Slide VI microfluidic channels were connected to the gravity-driven setup within a sterile hood. The hydrostatic chamber was then programmed to maintain a constant 40 mm fluid-height (392 Pa inlet pressure) to drive flow through the chip which had its outlet at atmospheric pressure through the dripping recirculation circuit. Laminar flow was assumed given the imposed pressure gradient and the rectangular design of the IBIDI device channel (length=17 mm, width=3.8 mm and height=0.4 mm). The Poiseuille equation was used to calculate the expected flow rate given a hydrostatic pressure difference *Q* = Δ*P/R*; where *Q* is the volumetric flow rate, Δ*P* is the hydrostatic pressure difference (392 Pa for a 40-mm media height) and *R* is the total resistance of the fluidic circuit (*R*_*Channel*_ + *R*_*tubing*_). For the IBIDI channel, which has a rectangular cross section, the fluidic resistance can be calculated as ^35,36,37^:

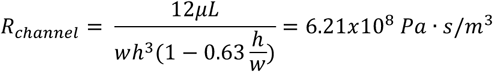

**Figure 8.**
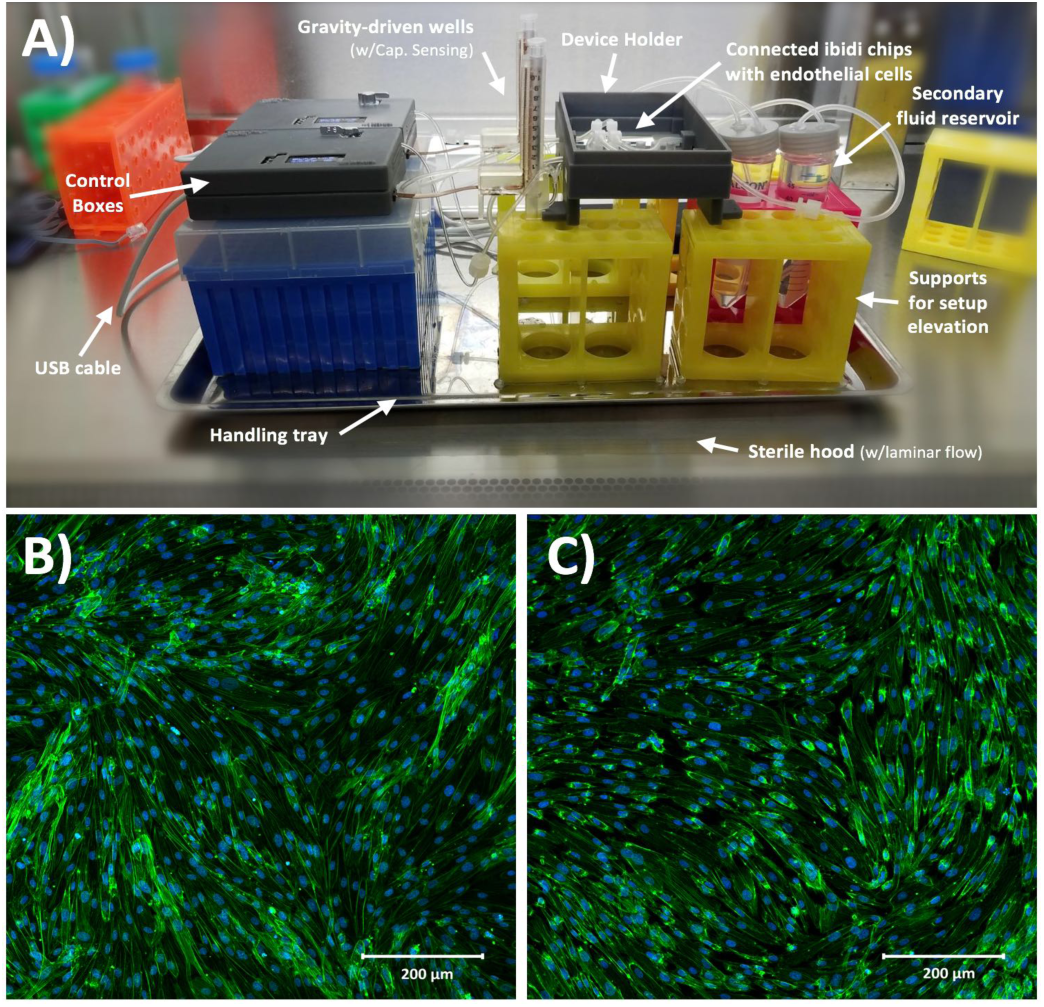
A) Closed-loop gravity-driven setup used for cell-culture experiment. B) Confocal microscopy of endothelial cells cultured under static conditions as control. C) Confocal microscopy of endothelial cells cultured under 4cm of closed-loop gravity-driven flow. Both cultures present with no observable differences, which suggests appropriate biocompatibility of the system.

Where *μ* is the dynamic viscosity of media which was assumed to be close to that of water (6.94x10^−4^ Pa·s at 37 °C), *L* is the length of the channel, *w* is the width and *h* is the height.

For the tubing, which has a circular cross-sectional area with inner diameter 1/32 inch and total length of 20 cm, the fluidic resistance can be calculated as^36^:

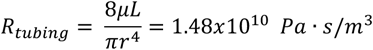

Where *r* is the inner radius of the tubing, *L* is the length of the tubing, *μ* is the dynamic viscosity of media which was assumed to be close to that of water (6.94x10^−4^ Pa-s at 37 °C). Using this information, we calculated the flow rate *Q* to be 2.64x10^−8^ m^3^/s or 1.59 mL/min at this specific height. The wall shear stress, *τ*, imposed on the endothelial cells under this flow rate can be calculated as^38^:

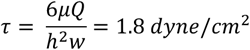

Here *τ* is in dyne/cm^2^, *μ* is in poise, *Q* is in cm^3^/s and *h* and *w* are in cm and are the height and width of the IBIDI channel respectively. After 24 hours of culture, cells were fixed with 4% PFA, stained with DAPI and Rhodamine and imaged under an EVOS inverted microscope (Life Technologies, Woburn MA, USA) with a 20x objective.

### II.c Statistical analysis

All validation experiments were conducted in triplicates. Error bars represent standard deviation in all figures. Calculations and plots were generated with Graphpad Prism 7 (GraphPad Software Inc.; La Jolla, USA).

## III. Results & discussion

### III.a Accurate capacitance fluid sensing and fluid tracking

The results of the initial characterization experiments used to assess the accuracy of capacitance readings and basic fluid level tracking capabilities of our sensing circuit are shown in Figure 4. Measured values of known capacitors directly connected to the C_sense_(+) and C_EXC_ terminals in our sensing circuit appear to match those of the E4981A capacitance meter within <0.1 pF error (Figure 4A). Readings from the capacitive sensing circuit also appear to be linearly related to those obtained from the calibrated reference meter suggesting appropriate implementation of the sensor at the PCB level. Figure 4B shows the aggregated results for three basic fluid-level tracking challenges using the fluid chamber with the proposed excitation-sensing-excitation inter-digitating capacitive sensor arrangement (ESE-ID) as seen in Figure 2C. The dashed red profile in Figure 4B is the known fluid-height as imposed by the calibrated syringe pump. Black data points refer to the averaged capacitive sensor output for the three replicates at each time point. The average standard deviation across all samples in this experiment was <250 μm when compared to the fluid height imposed by the calibrated syringe pump. Figure 4C shows both measured and estimated flow-rates in our gravity-driven pump as a function of set fluid height. Experimental measurements closely follow the theoretical values predicted by Poiseuille’s equation. Error bars are shown in green for all measurements in Figure 4. If further error reduction is required, averaging techniques and low-pass filtering may also be implemented in longer experimental time scales.

### III.b Capacitance readings depend on fluid conductivity

Figure 5A shows the results from the titration experiments conducted to characterize performance of our capacitive sensor based on fluid conductivity. Capacitance readings for upper (H_1_=30 mm) and lower (H_0_=10 mm) levels were smallest and closest together for deionized (DI) water and increased as electrolyte concentration increased to 1x PBS (conductivity @25°C = 1.6 S/m). Small steel rods of both lengths were also placed inside the monitored reservoir to assess signal in the presence of a known perfect conductor. Higher dilutions (0.5x to 0.0625x PBS) led to a range reduction and lower absolute capacitance, reaching a minimum in DI-water lacking solutes (conductivity @25°C = 0.055 μS/cm).

This behavior can be explained by the model shown in Figures 5B and 5C. In the case of water with dissolved solutes forming free ions (Figure 5B), a large number of sensing fringing paths become terminated near the fluid-reservoir boundary leading to higher capacitance measurements. Conversely, in the case of pure water (Figure 5C), fringing capacitance takes longer uninterrupted paths through the bulk of the fluid, generating smaller capacitance measurements. Figure 5D shows the simulated displacement flux density around the 1xPBS solution (ε_r,PBS_ = 80, σ_PBS_ = 1.45 S/m). The charge relaxation time of the 1xPBS solution is τ_PBS_ = ε_r,PBS_ ε_0_/σ_PBS_ = 488 ps, which corresponds to a break frequency f_PBS_ = 2.04 GHz. This break frequency is orders of magnitude higher than the 32kHz sensor excitation frequency used for the sensor.

Since f_PBS_ ≫ f_EXT_, the 1xPBS solution behaves like a conductor, accumulating free charges on its surface and increasing the apparent capacitance seen from the sensor electrodes. The induced surface charges terminate the flux density vectors, and therefore the flux density vectors are invisible inside the 1x PBS solution in Figure 5D. For the case of DI-water (ε_r,DI_ = 80, σ_DI_ = 5.5 μS/m), Figure 5C shows the field lines schematically, while Figure 5E shows the simulated flux density field. The charge relaxation time of the DI-water is τ_DI_ = ε_r,DI_ ε_0_/σ_DI_ = 129 μs, which corresponds to a break frequency f_DI_ = 7.76 kHz. Since f_DI_ < f_EXT_, the DI-water behaves like an insulator, which explains why the flux density vectors induced inside the DI-water follows the pattern shown in Figures 5C and 5E. These results show that calibration is required to adjust for potential differences in fluid conductivity. Further sections of this work make use of sensors calibrated for 1x PBS and culture media.

### III.c Accurate flow-rate tracking and calibration of external pneumatic micropumps

Figure S3 of the supplemental material shows the results of an extended series of experiments carried out to verify if accurate flow-rate calculations were achievable from analyzing fluid-height changes over time using our capacitive sensor. From these experiments, it was confirmed that a variety of flow-rate conditions can be inferred based on this sensor’s output and that these values correspond to the programmed settings in the calibrated syringe pump used to impose flow. Figure 6 shows the results of the characterization and calibration experiment of two open-loop pneumatic diaphragm micropumps (Figure 6A) feeding and extracting fluid from the same monitored reservoir at a nominal rate of 1 μL/s. Over the course of 40 min, a 0.2 mL decrease in fluid volume was observed in the monitored reservoir (Figure 6B), showing that the output pump flow-rate was slightly greater than the input pump flow rate despite being actuated at the same frequency.

Independent flow-rate characterization of each of these micropumps using our capacitive sensor revealed an 8.3% mismatch between the output and input micropump flow rates. This difference explains the observed decreasing height drift and is well within the 10-15% expected error usually reported for these type of pumping systems.^24^ After recalibration of the actuation frequency for both micropumps based on these readings, fluid-height drift appeared to be corrected for the same 40-min period as compared to the uncalibrated behavior. In general, long-term drift is expected to be present for a wide range of open-loop micropumps, due to back pressure, use-induced stress or solute deposition may sporadically change stroke volume. Thus, our sensor could be a valuable addition to open-channel microfluidic systems requiring accurate flow control, or as a new way to assess flow-rates and perform pump calibrations on demand.

### III.d Reliable closed-loop control of gravity driven pump

From available approaches for fluid handling in micro-and mesofluidic devices, gravity-driven systems have historically been considered among the most robust, simple and convenient to use.^13^ Reservoirs acting as gravity-driven pumps are relatively low-cost, can achieve a wide range of flow-rates, rarely lead to bubble stagnation and usually do not require external power to impose flow.^13^,^39^ However, traditional gravity-driven pumps (with vertically positioned reservoirs) can only produce unidirectional transient flows as the liquid level in the reservoir decreases.^13^,^16^,^22^ This situation leads to a time-dependent reduction in achievable flow rate proportional to the decline in hydrostatic pressure. This transient mode of operation is a key limitation of most gravity-driven systems, especially in long-term cell culture applications.^22^ Recent modifications of gravity-driven systems have been reported to provide nearly constant flow rates either through the use of horizontal reservoirs (setting a deterministic internal fluid height) or through the use of a large vertical reservoir (maintaining fluid heights nearly constant during limited operation times).^40^,^41^,^42^ Despite the advantages that these modifications may provide, most currently reported gravity-driven microfluidic devices still remain open-loop in nature, are cumbersome to continuously monitor and cannot deliver bidirectional, smooth, reconfigurable flow over long periods of time. Without closed-loop feedback control, the emptying time for an initial 30 mm fluid column at the hydrostatic chamber in connection with the used single-channel microfluidic chip was approximately 265 sec.

Figure 7 shows the typical closed-loop response provided by our gravity-driven microfluidic setup to a static (Figure 7A) and dynamic (Figure 7B) target. Both graphs show the aggregated height measurements for three experimental replicates as a function of time. Capacitive readouts were acquired every 10 ms, but were continuously averaged over 10 samples (T=100 ms) in both cases to feed the control algorithm. The maximum standard deviation across the entire 48-hr testing period for the constant target set point (ΔH=30 mm) was 0.65 mm. Figure 7B, shows the results obtained over a 40-min period using a dynamic pre-programmed set point. The constant, sine, triangular, saw tooth and step waveforms were all recognizable and accurately followed with less than 5% error. Overshooting decaying oscillations were observable at the high-frequency transitions for both step-like cycles, which is characteristic of many second order systems using closed-loop feedback control as it reflects the control loop dynamics.

In our augmented gravity-driven setup, fluid height appears to be a useful target variable allowing us to accurately control both pressure and flow-rate in this system. While these variables are commonly measured and used to control large-scale gravity-dominated fluidic systems (volumes > 10 L)^43^,^44^, this is the first time, to our knowledge, that capacitive sensing has been adapted to control such parameters in an open-well microfluidic system. Thus, we believe that previous limitations associated with sensing accuracy, miniaturization, ease of use and cost of implementation, may be addressed by our proposed sensor design.

### III.e Biocompatibility for continuous cell culture

Finally, Figure 8A shows the assembled and sterilized gravity driven setup used for our cell culture proof-of-concept. Optical analysis of both static (Figure 8B) and gravity-driven (Figure 8C) cultures confirmed viability. No differences in endothelial growth was observed in the test samples compared to the control, which suggests adequate biocompatibility of the designed fluidic circuit, as expected due to the use of inert materials in the system’s fluidic pathways. Given the low shear stress we exposed the cells to (1.8 dyn/cm^2^), we did not expect alignment in the direction of flow as such response for endothelial cells typically requires a shear >10 dyne/cm^2^ and in previous characterization of iPSC-derived endothelial cells a 20 dyne/cm^2^ shear stress was used.^45^

## Conclusions

Previous to this work, accurate measurement of fluid-height in open reservoirs for microfluidic devices had been considered a challenging task, especially in applications using total volumes below 1mL Traditional fluid-level sensing technologies such as mechanical floaters, displacers, bubblers, magnetic level gauges, magnetostrictive level transmitters and ultrasonic distance sensors are all currently inadequate or unreliable at the scale and operation of most micro-and mesofluidic devices.^46^ Furthermore, these limitations have been exacerbated by the fact that most previous approaches required some degree of fluid contact preventing their use in sterile biological applications. More recent options such as inline flow sensors and differential pressure transducers have become increasingly smaller and sensitive over the past years, and are now used in several microfluidic applications.^47,48,49^ Non-contact fluid-level sensing technologies; on the other hand, have only become popular in a limited number of large-scale industrial applications including monitoring and handling of large volumes of hazardous fluids. From direct visualization of fluid height (e.g. using optical monitoring through translucent windows), to time-of-flight (TOF) measurements (e.g. ultrasonic, laser and radar transmitters), and capacitive fluid sensing^50^; non-contact fluid-level sensors are now an increasingly common strategy to control macrofluidic systems where relevant height changes happen in the order of tens of centimeters. From these non-contact options, capacitive fluid sensing is perhaps one of the most attractive due to scalability and low cost. In the case of microfluidic applications, other groups have previously demonstrated the use of capacitive sensing for binary detection of droplets^51^ and droplet volume quantification.^17^,^52^ Furthermore, additional research groups have used this technology to develop contact-based in-line pressure sensors^52^, assess composition of liquids^53^ and even flow directionality sensors.^54^ These reported technologies demonstrate the general versatility and ease of integration of capacitive sensing; however, to our knowledge we are the first group to use this type of sensing technology to perform continuous measurements of fluid height changes in open micro-and mesofluidic reservoirs at submillimeter spatial resolution and millisecond temporal resolution to provide closed-loop control of pressure gradients and flow through a gravity driven system.

Though limited in some ways (e.g. dependent on fluid properties and requiring non-conducting fluid chambers we believe the design principles presented here can be used to adapt such sensors to a wide range of open-channel microfluidic devices. In particular, we believe this technology may allow for continuous fluid height monitoring in high throughput microtiter plates and cell culture platforms with the help of robotic liquid handlers, or even in the monitoring of gravity driven chambers to measure permeability of small porous materials such as scaffolds and hydrogels.

## Acknowledgements

The authors would like to thank the microfluidics community who have provided both direct and indirect input for the design specifications the presented sensor and system. This work was funded under the Defense Advanced Projects Agency (DARPA) Microphysiological Systems Program (W911NF-12-2-0039).

## Supplementary Information

**Figure S1.**
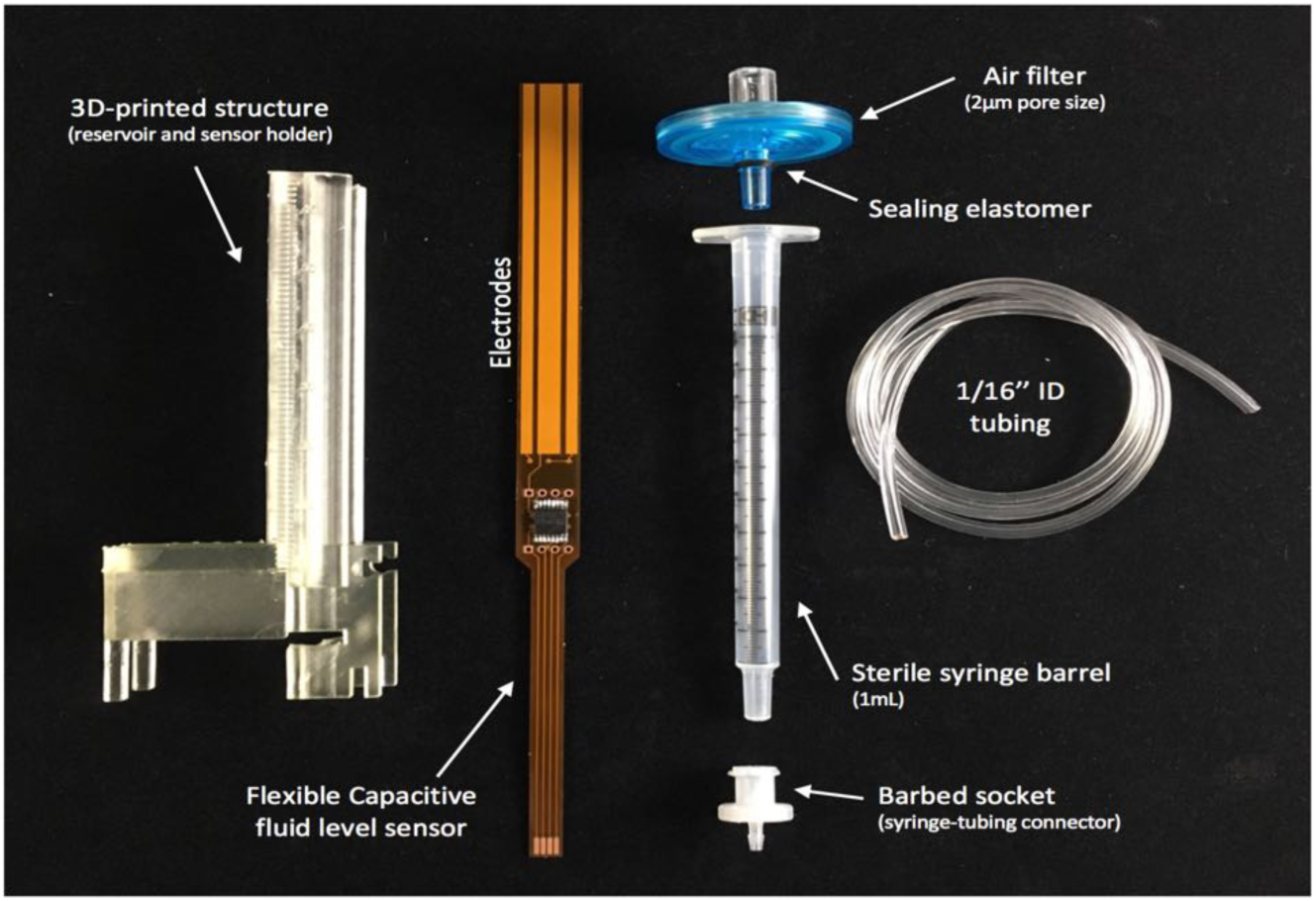
Detail on individual components used for the assembly of the monitored hydrostatic chamber used for the feedback-controlled gravity-driven setup. For assembly, the syringe barrel, barbed socket, tubbing and filter are connected together to be sterilized and then assembled onto the 3D-printed structure. The capacitive sensor is inserted into a thin slot at the back of the 3D-printed structure that positions the sensor in close proximity to the syringe barrel and thus the fluid. The sensor can be permanently attached to the 3D-printed structure with resin or adhesive to avoid mechanical induced drift.

**Figure S2.**
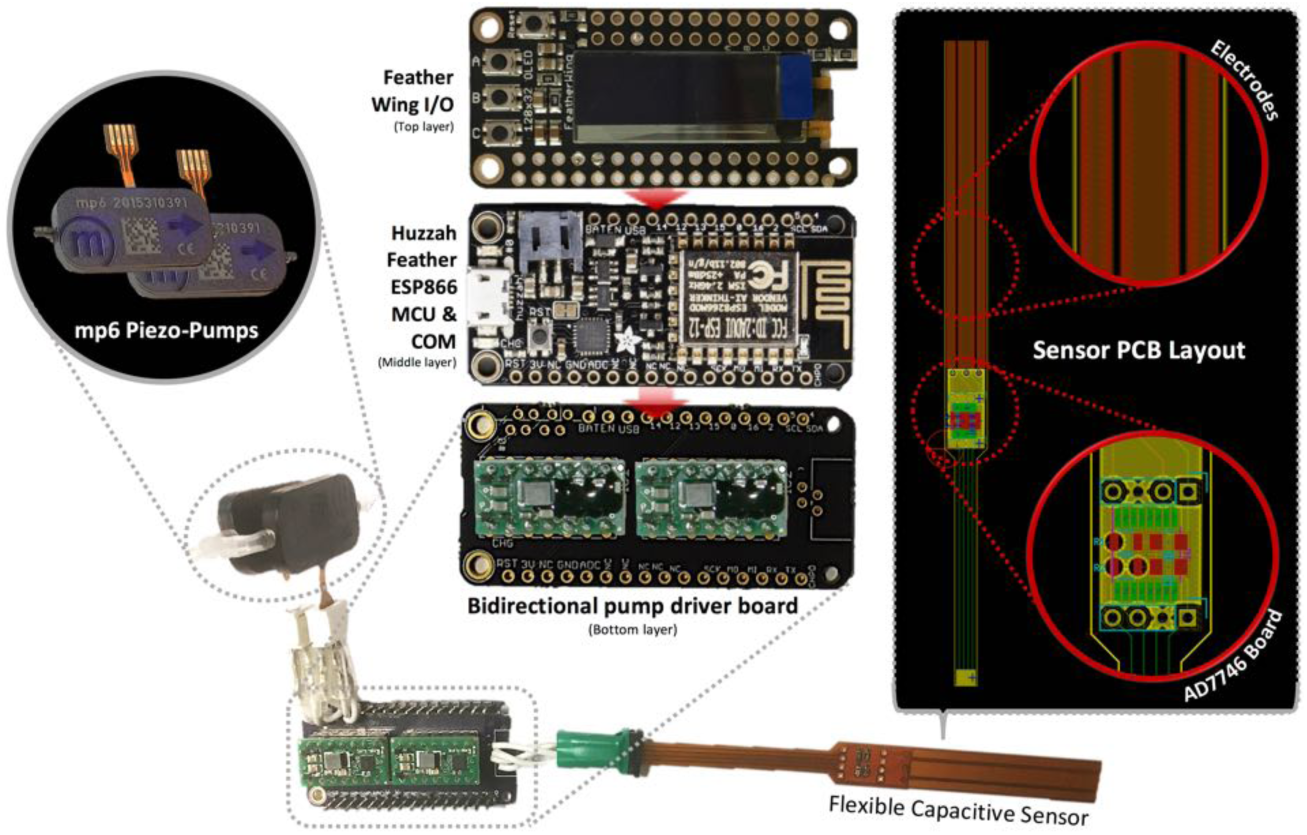
Individual components of control hardware. Two mp6 piezoelectric pumps (top left) were connected to a bidirectional pump driver shield (bottom). This pump driver shield was controlled by a stackable Huzzah Feather ESP866 MCU & COM board containing the microcontroller unit. An OLED screen, as well as input keys can be seen in the Feather Wing I/O shield, which also interphase to the micro-controller board. The flexible capacitive sensor can be seen connected to the pump driver shield to interfase with the micro-controller (bottom). A close-up of the sensor PCB layout with details of the electrodes and AD7746 board is also shown (right).

**Figure S3.**
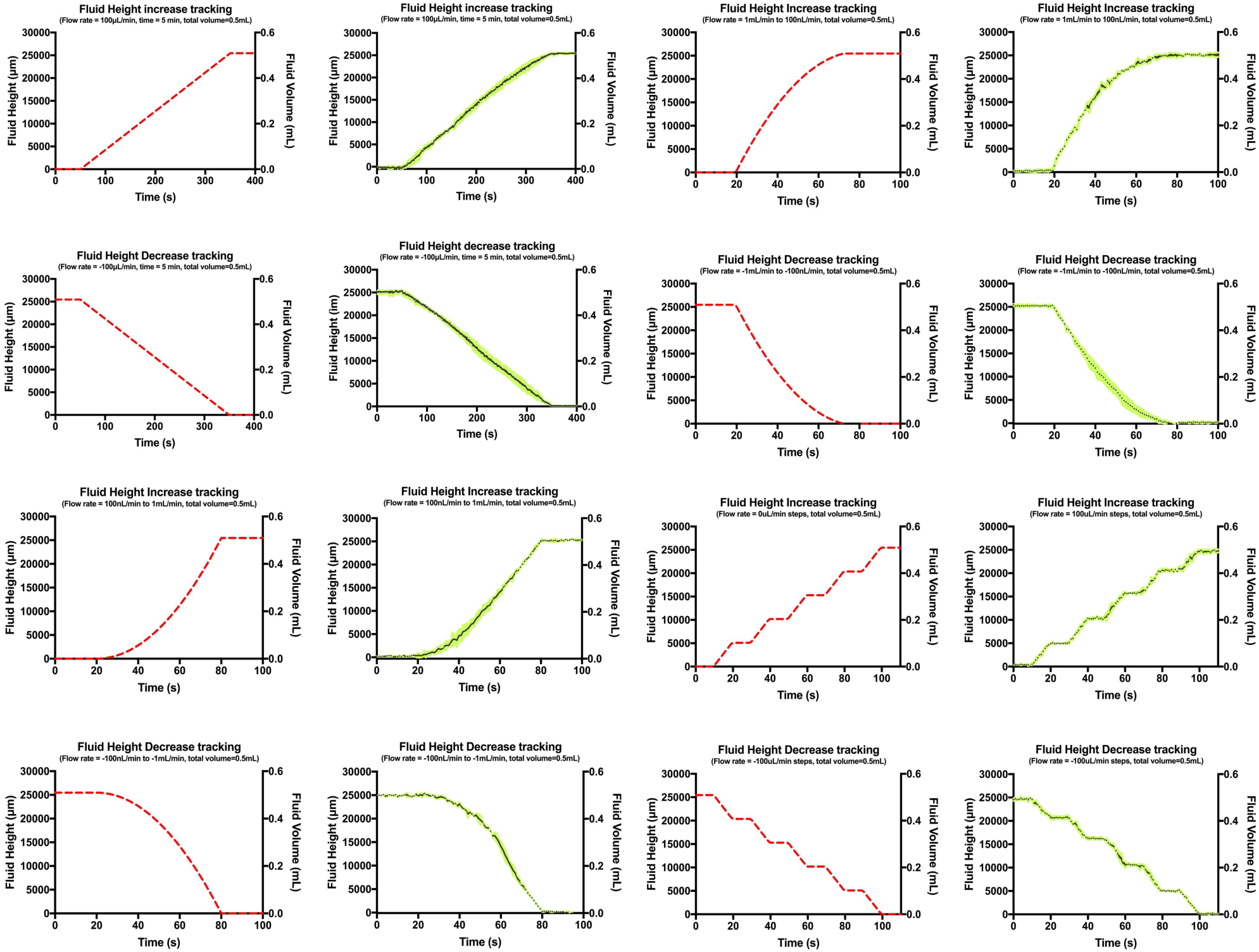
Results of fluid height tracking with constant, and increasing/decreasing ramp and step flow-rate profiles. Programmed flow-rate settings for each experiment are shown below graph titles. Red dashed profiles denote the predicted fluid height and volume change over time as programmed in the calibrated syringe pump. Black points denote the average of three replicates following the same experimental conditions at each time point, while green error bars refer to the standard deviation of those samples. Inflow and outflow was tested for constant 100ul/min flow-rate, as well as for increasing and decreasing ramps (ranging from 100nl/min to 1mL/min). Finally, inflow and outflow experiments were also conducted for a step-like profile with 100ul/min amplitude. Flow rates were calculated every 10s and compared with flow rates know from simulations. Constant and dynamic additions or extractions of fluid generated linear fluid height increase and decrease profiles that were similar for both simulations and experimental results. Increasing and decreasing ramp profiles generated second order profiles similar to those predicted by simulations. Similar results were found for the step flow profiles.

